# Estimating and correcting index hopping misassignments in single-cell RNA-seq data

**DOI:** 10.1101/2024.10.21.619353

**Authors:** Lingling Miao, Loren Collado, Savannah Barkdull, Yoshine Saito, Jay-Hyun Jo, Jungmin Han, Stefania Dell’Orso, Michael C. Kelly, Heidi H. Kong, Isaac Brownell

**Affiliations:** Dermatology Branch, NIAMS, NIH, Bethesda, Maryland, USA; Genomic Technology Section, NIAMS, NIH, Bethesda, Maryland, USA; Single Cell Analysis Facility, Cancer Research Technology Program, Frederick National Laboratory, Bethesda, Maryland, USA

**Keywords:** Single-cell RNA sequencing, barcoding, multiplexing, index hopping, read misassignment, Merkel cell, dermal papilla, hematopoietic cell

## Abstract

**Background:** Index hopping causes read assignment errors in data from multiplexed sequencing libraries. This issue has become more prevalent with the widespread use of high-capacity sequencers and highly multiplexed single-cell RNA sequencing (scRNA-seq).

**Results:** We conducted deep, plate-based scRNA-seq on a mixed population of mouse skin cells. Analysis of transcriptomes from 1152 cells identified four distinct cell types. To estimate the error rate in sample assignment due to index hopping, we employed differential expression analysis to identify signature genes that were highly and specifically expressed in each cell type. We quantified the proportion of misassigned reads by examining the detection rates of signature genes in other cell types. Remarkably, regardless of gene expression levels, we estimated that 0.65% of reads per gene were assigned to incorrect cell across our data. To computationally compensate for index hopping, we developed a simple correction method wherein, for each gene, 0.65% of the library’s average expression level was subtracted from the expression in each cell. This correction had notable effects on transcriptome analyses, including increased cell-cell clustering distance and alterations in intermediate state assignments of cell differentiation.

**Conclusions:** Index hopping misassignments are measurable and can impact the experimental interpretation of sequencing results. We devised a straightforward method to estimate and correct for the index hopping rate by quantifying misassigned genes in distinct cell types within an scRNA-seq library. This approach can be applied to any barcoded, multiplexed scRNA-seq library containing cells with distinct expression profiles, allowing for correction of the expression matrix before conducting biological analysis.

## BACKGROUND

The advent of next-generation sequencing has revolutionized genomic research, with contemporary sequencers allowing cost-effective sequencing of large-scale multiplexed libraries [1]. Multiplexing is a common strategy for sequencing multiple samples that involves adding a unique short DNA sequence (index) to each fragment in a sample and pooling samples together for sequencing. However, a notable limitation of this approach is index misassignment, where sequence reads are erroneously paired with an index from another sample. Index misassignment can occur at various stages during sequencing including reagent contamination, sample handling, PCR amplification, patterned flow cell "spread-of-signal", and bioinformatic errors [2, 3]. Among these mechanisms, a major contributor to index hopping (also known as index swapping or index switching), is incomplete library cleanup which allows free-floating index primers to randomly amplify DNA fragments from multiple samples within a pooled library during sequencing [4]. This phenomenon leads to data contamination through index misassignment. The impact of index hopping has grown significantly as high capacity modern sequencing technologies increasingly rely on multiplexing samples. This issue is particularly evident in single-cell RNA sequencing (scRNA-seq) studies that involve pooling large numbers of cDNA libraries for sequencing [4–7].

Index hopping poses a challenge in scRNA-seq, impacting both droplet and plate-based methods [4–7]. Recent investigations analyzing multiplexed droplet-based scRNA-seq datasets estimated sample index hopping probabilities ranging from 0.3% to 0.9% [4]. Plate-based scRNA-seq studies utilizing the "i5+i7" dual-index multiplexing strategy reported more variable and higher index hopping rates, ranging from 0.25% to 10% on Hiseq platforms [5–7]. The index hopping rate varies across different Illumina platforms, with advanced Hiseq systems, such as Hiseq 3000, which employ patterned flow cells and exclusion amplification chemistry, exhibiting higher index hopping levels compared to earlier platforms like Hiseq 2500 [3–5, 8, 9]. Illumina recommends reducing free index primers or the use of unique dual indexes as experimental approaches to mitigate index hopping when using dual-index multiplexing strategies [10]. However, unique dual indexing may not be practical with large samples due to the limitation in barcode numbers, especially in high-volume scRNA-seq systems. In addition, unique indexing is not a complete solution as index hopping has been observed in droplet-based scRNA-seq and multiplexed long-read sequencing which use unique sample barcodes [4, 5, 11]. A few recent studies have explored using computational algorithm to estimate and correct the contamination effect of index hopping in plated-based and droplet-based scRNA-seq data [4–6]. However, their approach requires knowing the ground truth of non-multiplexed sequencing.

Given the limitations of existing strategies, there is a need for an objective method to estimate and correct for index hopping in scRNA-seq data. In this study, we performed an in-depth analysis of plate-based scRNA-seq data obtained from mouse skin cells. We developed a simplified method that leverages distinct expression profiles among cells to estimate the index hopping rate and implemented a correction approach to adjust for the contamination caused by index hopping. We further demonstrate the impact of correcting for index hopping before conducting biological analysis of scRNA-seq data. Our findings contribute to advancing the understanding of index hopping and provide a practical solution for its estimation and correction in scRNA-seq data analysis.

## METHODS

### Animals

*Sox2^GFP/+^* [12] and *Gfi1^GFP/+^* [13] mice were derived from breeders purchased from Jax (#017592 and #016162, respectively). Mice were housed and bred on an outcrossed Swiss Webster background in a pathogen-free facility at NIH, Bethesda, MD. Genotyping of mice was performed by allele-specific PCR on DNA extracted from tail tissue. All experiments were performed in accordance with institutional guidelines according to Institutional Animal Care and Use Committee-approved protocols.

### Cell dissociation and sorting

Postnatal day 0 (P0) *Sox2^GFP/+^* and *Gfi1^GFP/+^* mice were euthanized using CO2 inhalation followed by decapitation, and the skin was harvested, removing the subcutaneous tissue. For embryonic day 18.5 (E18.5) cells, the pregnant *Sox2^GFP/+^*female mice were euthanized using CO2 inhalation followed by cervical dislocation, and the embryos were harvested and euthanized. The skin was harvested from the embryos. The harvested E18.5 or P0 skin was floated in dispase (Corning, #354235) at room temperature for 1 hour to separate the epidermal sheet. The epidermis was chopped into small pieces, and the epidermal cells were dissociated in 0.05% Trypsin at 37 °C for 15 minutes, followed by the addition of DMEM containing 10% FBS. The cells were then sequentially filtered through 70 µm and 40 µm cell strainers. Subsequently, the cells were centrifuged at 300 x g for 5 minutes, and the supernatant was discarded. The cell pellets were resuspended and processed for cell sorting using the Beckman MoFlo Astrios or BD influx cell sorter.

DAPI was added immediately before cell sorting, and single DAPI^-^GFP^+^ cells were sorted into 96-well plates pre-loaded with lysis buffer (lysis buffer prepared following the smart- seq2 method [14]). Single DAPI^-^GFP^-^ cells from P0 epidermis were sorted as controls.

For embryonic day 16.5 (E16.5) and embryonic day 17.5 (E17.5) cells, pregnant *Sox2^GFP/+^*(E16.5) and *Gfi1^GFP/+^* (E16.5 and E17.5) female mice were euthanized using CO2 inhalation followed by cervical dislocation, and the embryos were subsequently harvested and euthanized. The skin from the embryos was then harvested, chopped into small pieces, and treated with 0.05% Trypsin at 37 °C for 15 minutes. This was followed by the same cell dissociation procedure used for P0 and E18.5 skin, without separating the epidermis.

Postnatal day 6 (P6) *Sox2^GFP/+^*and *Gfi1^GFP/+^* mice were euthanized using CO2 inhalation followed by decapitation. The harvested skin was then incubated in 0.15% trypsin at 37°C for 30 minutes. The epidermis was obtained by scraping with a scalpel in 0.05% trypsin. Epidermal cells were then processed using the same method employed for neonatal epidermis.

### scRNA-seq

Libraries from isolated single cells were generated following the Smart-seq2 protocol [14] with the following modifications. Prior to tagmentation, the concentration of single-cell cDNA libraries was determined using the Quant-iT PicoGreen dsDNA Assay (Thermo Fisher Scientific). Subsequently, the cDNA libraries were relocated to new plates with normalized concentration. Libraries from each set of 384 cells with unique barcode combinations were pooled together, resulting in each pooled library containing cells from all ages and both reporters. Sequencing was performed on the Hiseq3000 sequencer (Illumina), resulting in three pooled libraries. Additionally, the second pooled libraries underwent repeated sequencing on the Hiseq2500 sequencer (Illumina), achieving a sequencing depth comparable to that of the Hiseq3000. The sequencing generated paired-end reads of either 125bp or 100bp. Subsequently, reads of 125bp were trimmed to 100bp for further processing. The sequencing reads were mapped to the mouse mm10 reference genome using STAR 2.5.2b, and gene expression was quantified using RSEM 1.3.0. The gene expression matrix with transcripts per million (TPM) values was used for further analysis. Log2-scaled TPM values (log2(TPM+1)) were used in all figures where TPM was shown, unless otherwise specified (log2TPM in Figure 5G, Figure 5H, and Figure S6). For non-self genes in MCs, their average detection ratios in MCs are considered as the index hopping rate.

### Cell biology analysis

Cell-to-cell distance was analyzed using the R package ‘sincell’ based on Spearman correlation [15]. Cell clustering was analyzed using Seurat 2.3 [16]. Cell trajectory was analyzed using Monocle 2 [17, 18].

### Statistical analysis

For comparisons of gene detection in single cells between two different conditions (such as before vs. after correction, Hiseq2500 vs. Hiseq3000), paired t-test was applied.

## RESULTS

### Index hopping rates can be estimated by the frequency of non-self genes in scRNA-seq data

We performed plate-based scRNA-seq on FACS sorted dissociated mouse skin cells. The skin is a complex tissue that encompasses multiple cell types including, but not limited to, Merkel cells, dermal papilla fibroblasts, hematopoietic cells, and epithelial keratinocytes (Fig. S1). The transcription factor SRY-Box Transcription Factor 2 (SOX2) is expressed in Merkel cells and dermal papilla cells [19], whereas Growth Factor Independence 1 (GFI1) is expressed in Merkel cells and hematopoietic cells [13, 20].

We sorted single cells expressing GFP from the skin of *Sox2^GFP/+^* or *Gfi1^GFP/+^* reporter mice at various developmental stages, along with GFP-negative epidermal cells expected to be keratinocytes (Fig. 1A). Single cells were sorted into lysis buffer in 96 well plates, and the Smart-seq2 method was used to generate cDNA from these sorted individual cells. Cleaved cDNA was tagged with dual index adaptors prior to being pooled into three 384-sample sequencing libraries (Fig. 2A, S2A-C). On average, each cell yielded ∼3 million uniquely mapped reads, allowing for the detection of approximately 6000 genes per cell. By employing Seurat clustering, we grouped the individual cells into four clusters based on their transcriptome similarities (Fig. 1B).

**Figure 1.**
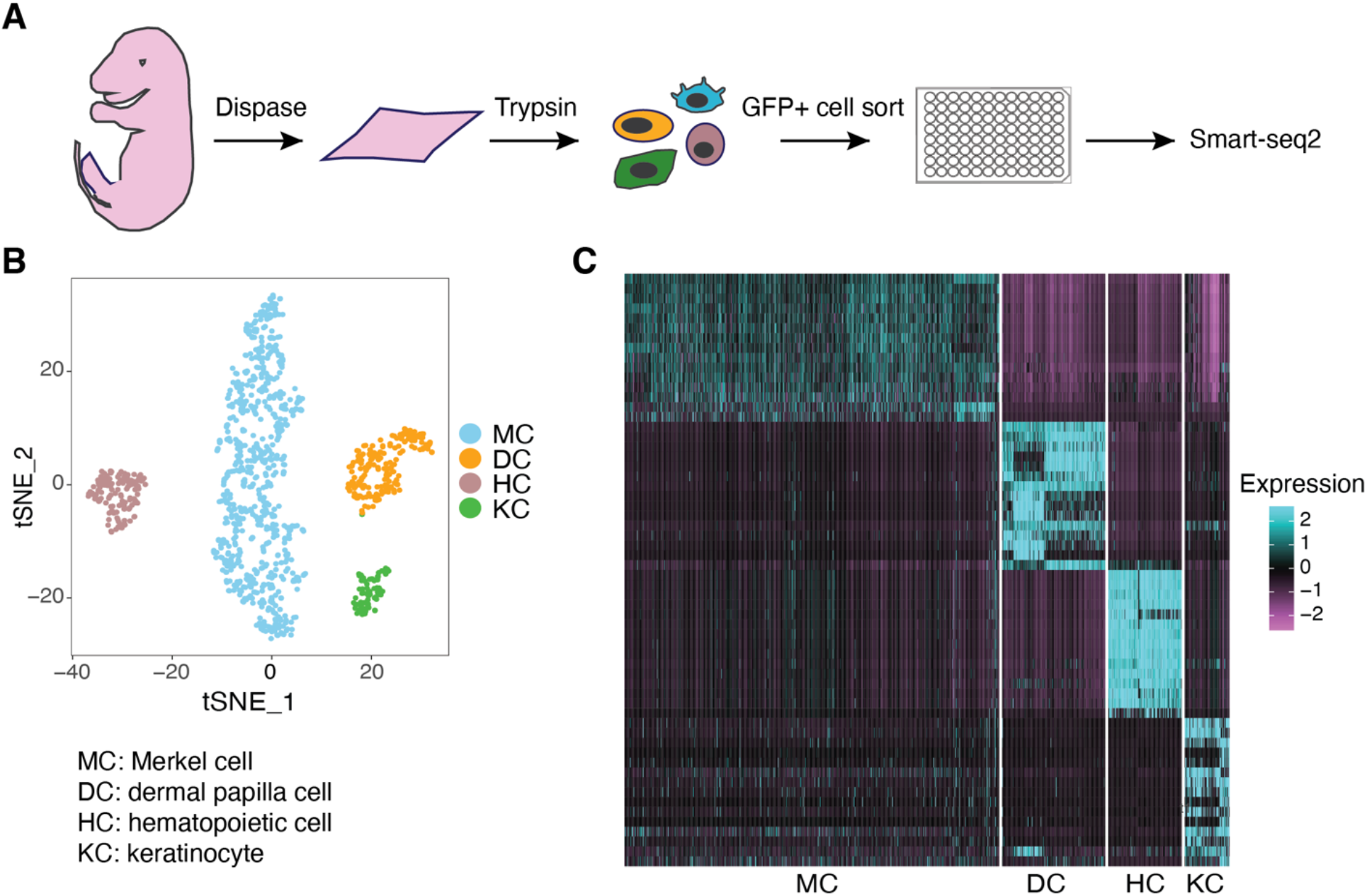
scRNA-seq identifies different skin cell types. **(A)** Overview of scRNA-seq experimental procedure. **(B)** tSNE plot showing clustering of the gene expression profiles of single skin cells. **(C)** Heatmap showing highly differentially expressed genes (marker genes) for each cell cluster identified by Seurat clustering.

**Figure 2.**
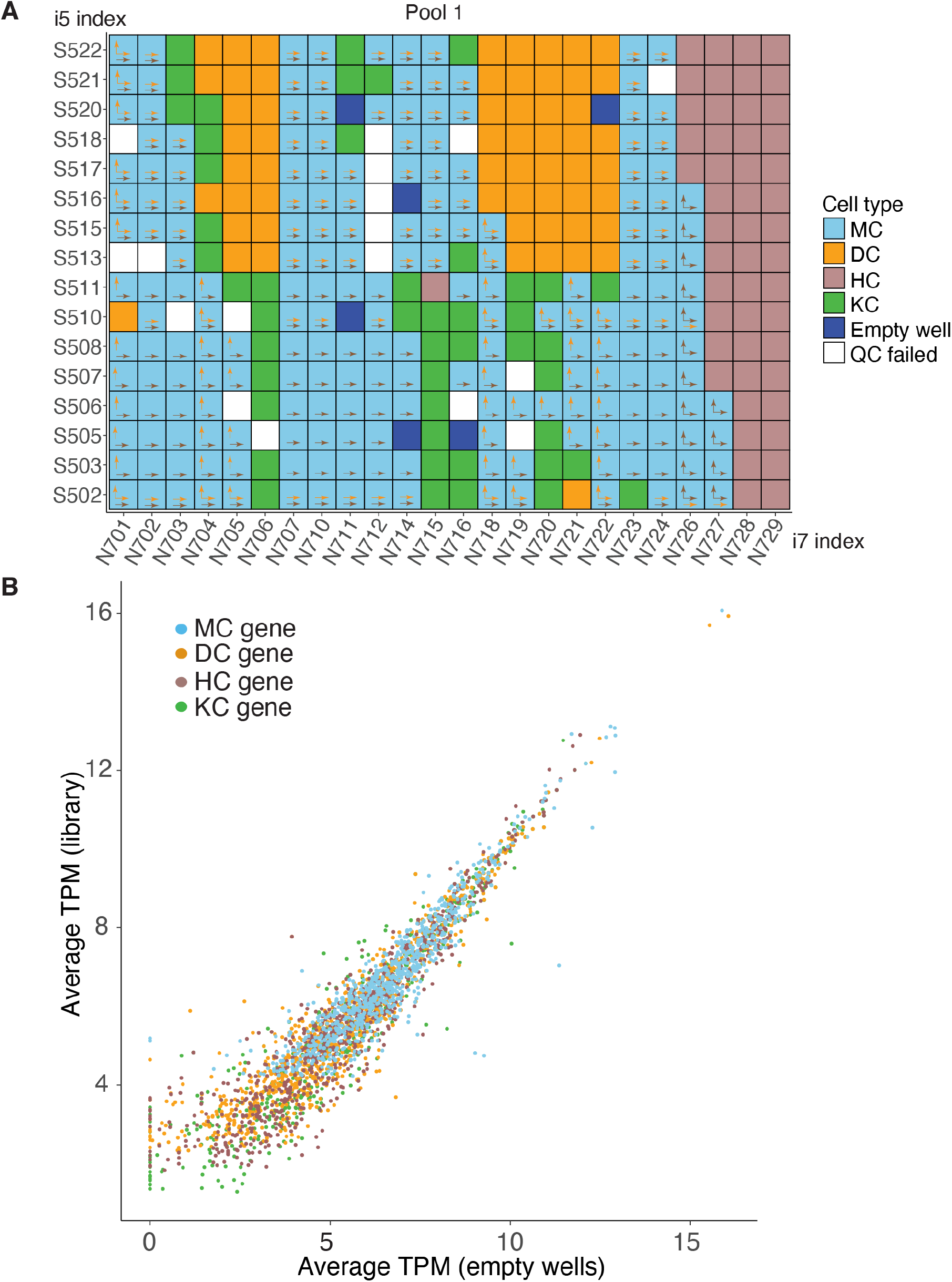
Index hopping can be detected when occurring between differing cell types. **(A)** Overview of the indexing used for 384 single-cell cDNA libraries in the first pooled sequencing library. Cell types together with empty wells and quality control failed wells are color-coded. Arrows show for each individual Merkel cells (MCs), the potential index hopping with dermal cells (DCs, orange arrow) or hematopoietic cells (HCs, brown arrow). Vertical arrows show i7 index hopping, and horizontal arrows show i5 index hopping. **(B)** Scatter plot showing the correlation of average TPM of cell type marker genes in empty wells and the entire library, with values transformed using a log2 scale (log2(average TPM +1). Color labels marker genes for the 4 cell types.

Furthermore, by identifying highly differentially expressed genes (marker genes) within these four clusters, we were able to distinguish them as four distinct cell types: Merkel cells (MCs), dermal papilla cells (DCs), hematopoietic cells (HCs), and keratinocytes (KCs) (Fig. 1C). As the pooled libraries included all four cell types, index hopping had the potential to misassign reads from one cell type to another based on their sharing of i7 or i5 indexes (Fig. 2A, S2A-B).

Index hopping occurs when cDNA from one cell is indexed erroneously with another cell’s barcodes. After library pooling, unincorporated free index primers associated with a row or column on the barcoding plate can potentially be added to cDNA from another row/column creating the risk for index hopping and misassignments of reads. Detection of such an event is difficult when the two wells contain similar transcriptomes. However, if the two cells are of different lineages and have distinct gene expression, index hopping contamination can be measured by detecting non-self genes (transcript not expressed in that cell type) in a specific cell or from an empty well in plated-based sequencing libraries. To identify non-self genes, we used the differentially expressed marker genes for each of the four cell types in our library (Fig. 1C). We were able to detect marker genes for each of the four cell types labeled with indexes for empty wells. We then calculated the average library expression levels for all genes by dividing the total TPM for each gene in the library by the number of cells in the library. Similarly, we calculated the average detection levels for each gene in empty wells by dividing the total TPM from the empty wells by the number of empty wells. Comparing the average gene detection levels in the library and in empty wells, we observed a positive correlation (Fig. 2B, S2D), indicating that misassignments were likely a result of a stochastic process like index hopping rather than technical errors in sample preparation.

To assess index hopping in our library, we quantified non-self gene detection in the four cell types. Considering the location of single cell samples within the multi-well plates and the application of dual index primer sets in our libraries, there was the potential for dual index hopping (misassignment can happen by either i5 or i7 hopping), single index hopping (misassignment can happen only by i5 or i7 hopping), and no potential index hopping between DCs and MCs, while dual index hopping and i5 index hopping were possible between HC and MC (Fig. 2A, S2A-B). To estimate the rate of read misassignments, we calculated the following: Firstly, we computed the average TPM of a gene in MCs by dividing the total TPM for that gene in MCs by the number of MCs (gene A as an example, 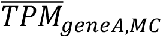 The data were shown on a log2 scale 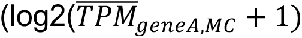.

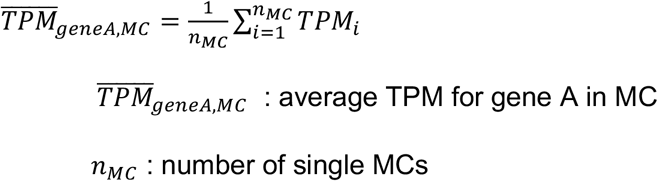

Then, to determine the ratio of reads assigned to individual MCs (detection ratio) for each gene (gene A as an example, 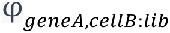), we divided the TPM of each gene in single MC by its average TPM in the library.

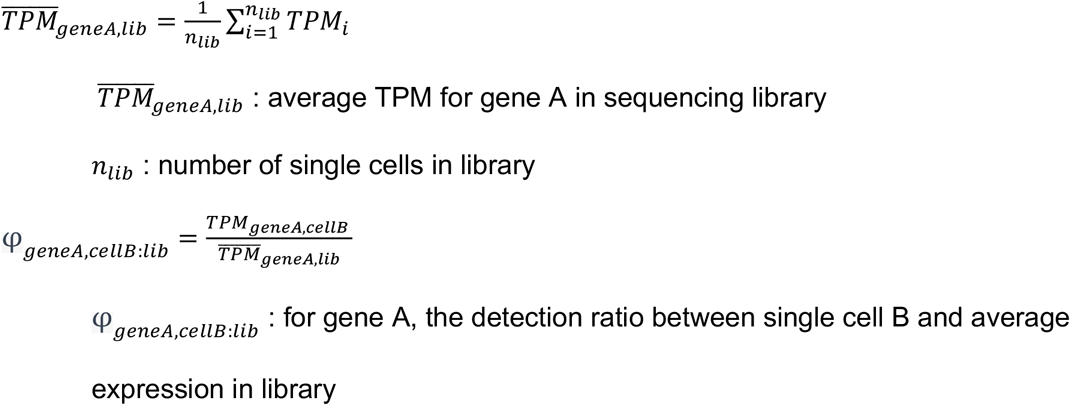

And we were able to calculate the average detection ratio of each gene between MCs and the entire library (gene A as an example,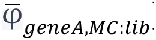).

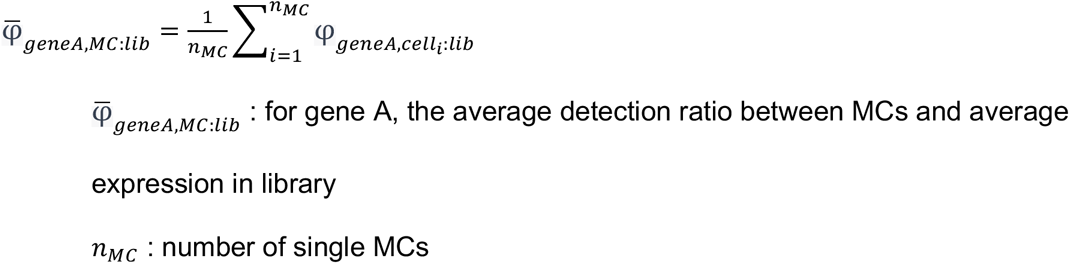

If index hopping played a significant role in misassignment, the detection ratio of non-self genes in MCs would correspond to the index hopping rate. Notably, the detection ratio of top DC genes in MCs was higher in MCs with dual index hopping from DCs compared to those with single index hopping from DCs. Conversely, when no index hopping occurred between MC and DC, the detection ratio of DC genes in MCs was nearly zero (Fig. 3A-B, S3A-B). Similarly, the detection ratio of top HC genes in MCs with dual index hopping from HCs was higher than those with i5 index hopping from HCs (Fig. 3C, D). Furthermore, the detection ratio of DC and HC genes in MCs exhibited a positive correlation with the number of index-hopped DCs (Fig. S3C) and HCs (Fig. S3D), respectively.

**Figure 3.**
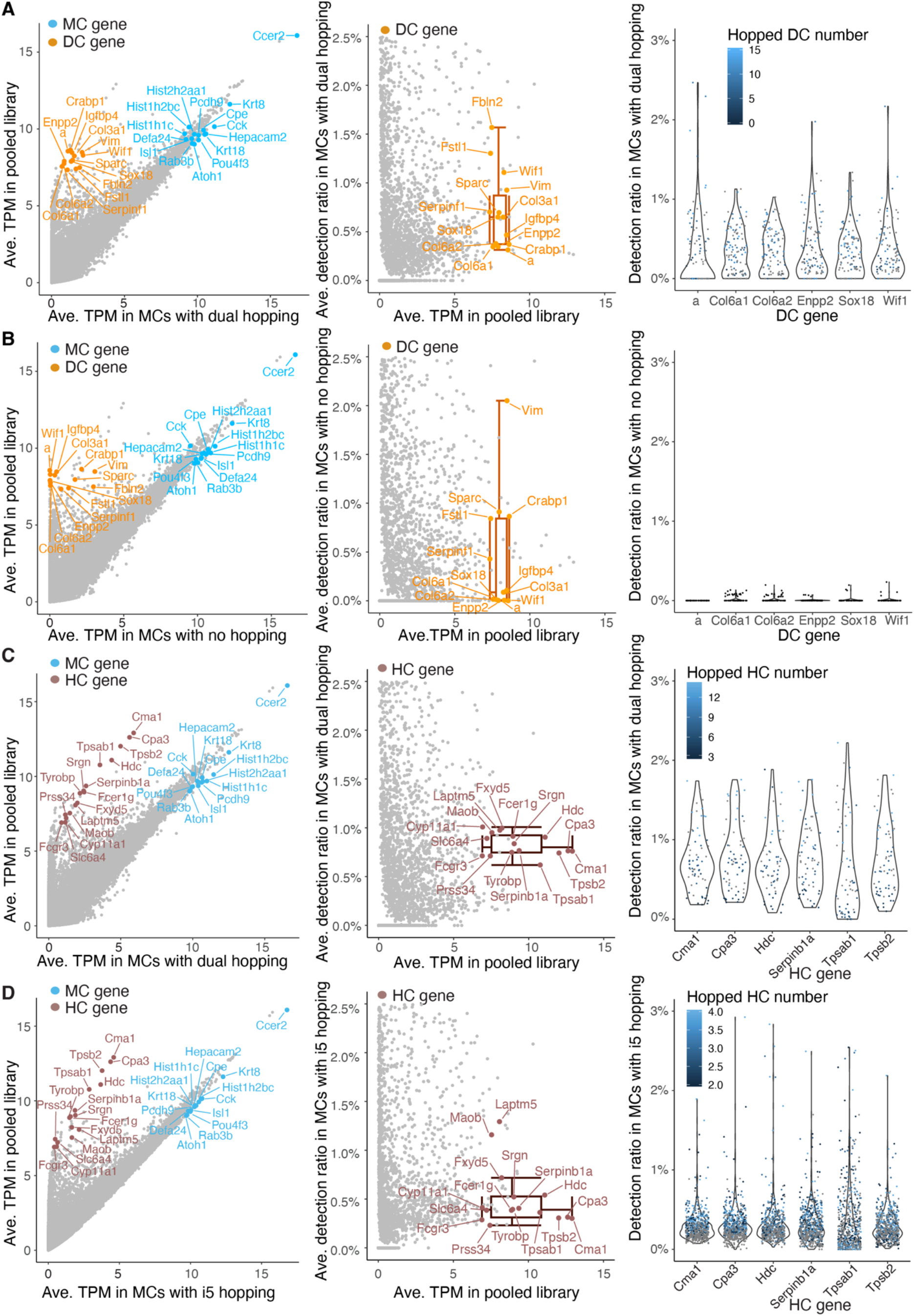
Non-self marker genes are detected at different levels in MCs with dual, single and no index hopping to other cell types. **(A)** and **(B)** MCs which had dual (A) and no (B) index hopping potential from DCs: Left, scatter plot showing the average TPM for individual genes in single MCs correlated with their average TPM in the library. TPMs are displayed on a log2 scale (log2 (average TPM + 1)). Middle, scatter plot showing the average detection ratios of DC genes in single MCs. Colored labels indicate the top marker genes for different cell types. Box plots show the median and interquartile range ± SD of average detection ratio for color-labeled genes. Ave., average. Right, violin plot showing the detection ratio of the top 6 DC marker genes in individual MCs **(C)** and **(D)** MCs which had dual (C) and i5 (D) index hopping potential from HCs: Left, scatter plot showing the average TPM in single MCs correlated with their average TPM in the library. TPMs are displayed on a log2 scale (log2 (average TPM + 1)). Middle, scatter plot showing the average detection ratio of HC genes in single MCs. Colored labels indicate the top marker genes for different cell types. Box plots show the median and interquartile range ± SD of average detection ratio for color-labeled genes. Ave., average. Right, violin plot showing the detection ratio of the top 6 HC marker genes in individual MCs.

### Adjusting gene expression levels by the estimated index hopping rate compensates for misassigned reads

Having observed that index hopping was relatively consistent, we sought to correct for its effect. A group of highly-expressed HC and DC marker genes with no expected expression in MCs was detected in MCs at levels consistent with their average TPM in the library (Fig. 4A). We referred to this distinctive group of genes with their misassigned reads in MCs as the “side population”. The detection ratios of these genes in MCs 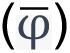 were consistently below 1%, regardless of their varying expression levels in the libraries (Fig. 4B). Notably, there were other highly-expressed HC and DC marker genes that appeared to have native expression in MC as indicated by their detection ratios being greater than 1%. These non-specific marker genes clustered away from the side population genes (Fig. 4A-B). To estimate average index hopping rates in MC, we employed a methodology inspired by the Rank Ordering of Super-Enhancers (ROSE) tool for super-enhancer identification from ChIP-seq data [21]. We first ranked and plotted all genes increasingly by the reciprocal of their average gene detection ratio 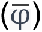 in MCs. This plot demonstrated a distinct inflection point where the reciprocal of the detection ratio in MCs began increasing rapidly. To geometrically define this point, we rescaled the data so that the x and y axes ranged from 0 to 1. Next, we determined where a tangent to the rank curve had a slope of 1. The gene detection ratio at this point was calculated to be 0.65%, which we used as the estimated index hopping rate in our sequencing library. Genes ranked below this threshold were classified as predominantly MC-expressed genes, whereas genes ranked above this threshold were considered as genes whose detection in MCs was due, in part, to index hopping misassignment (Fig. 4C). This classification was validated by identifying the rank order position of the differentially expressed marker genes for each cell type (Fig. S4A-D).

**Figure 4.**
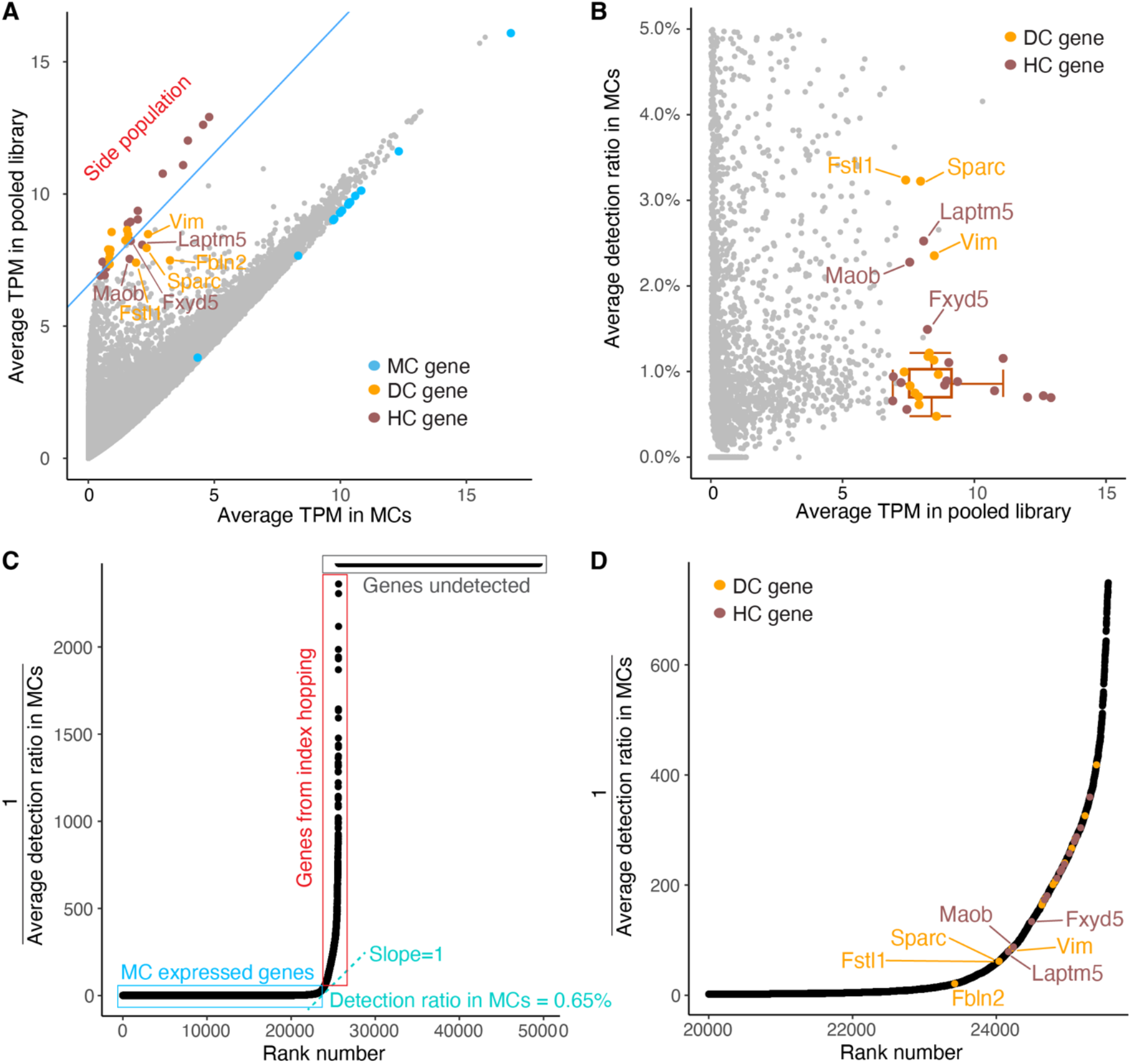
Index hopping rate can be estimated using non-self genes. **(A)** Scatter plot showing the average TPM of top HC and DC genes in single MCs correlated to their average TPM in library. Blue line separates highly-expressed HC and DC “side” population genes where the level in MCs is proportional to their average TPM in the library. TPMs are displayed on a log2 scale (log2 (average TPM + 1)). Colors indicate the top MC, HC and DC marker genes. **(B)** Scatter plot showing the average gene detection ratios in single MCs. Box plots show median and interquartile range ± SD of average detection ratio in MCs for side population genes. Colors indicate the top HC and DC marker genes. **(C)** Rank ordered scatter plot showing distribution of gene detection ratios in MCs across all genes from mm10 genome annotated by Ensembl. Blue box indicates the MC expressed genes, red box indicates genes presumptively detected due to index hopping, grey box indicates genes with no detection in MCs. The cyan tangent line with slope = 1 identifies the inflection point. **(D)** Focused rank ordered scatter plot showing genes near the inflection point from plot in (C).

**Figure 5.**
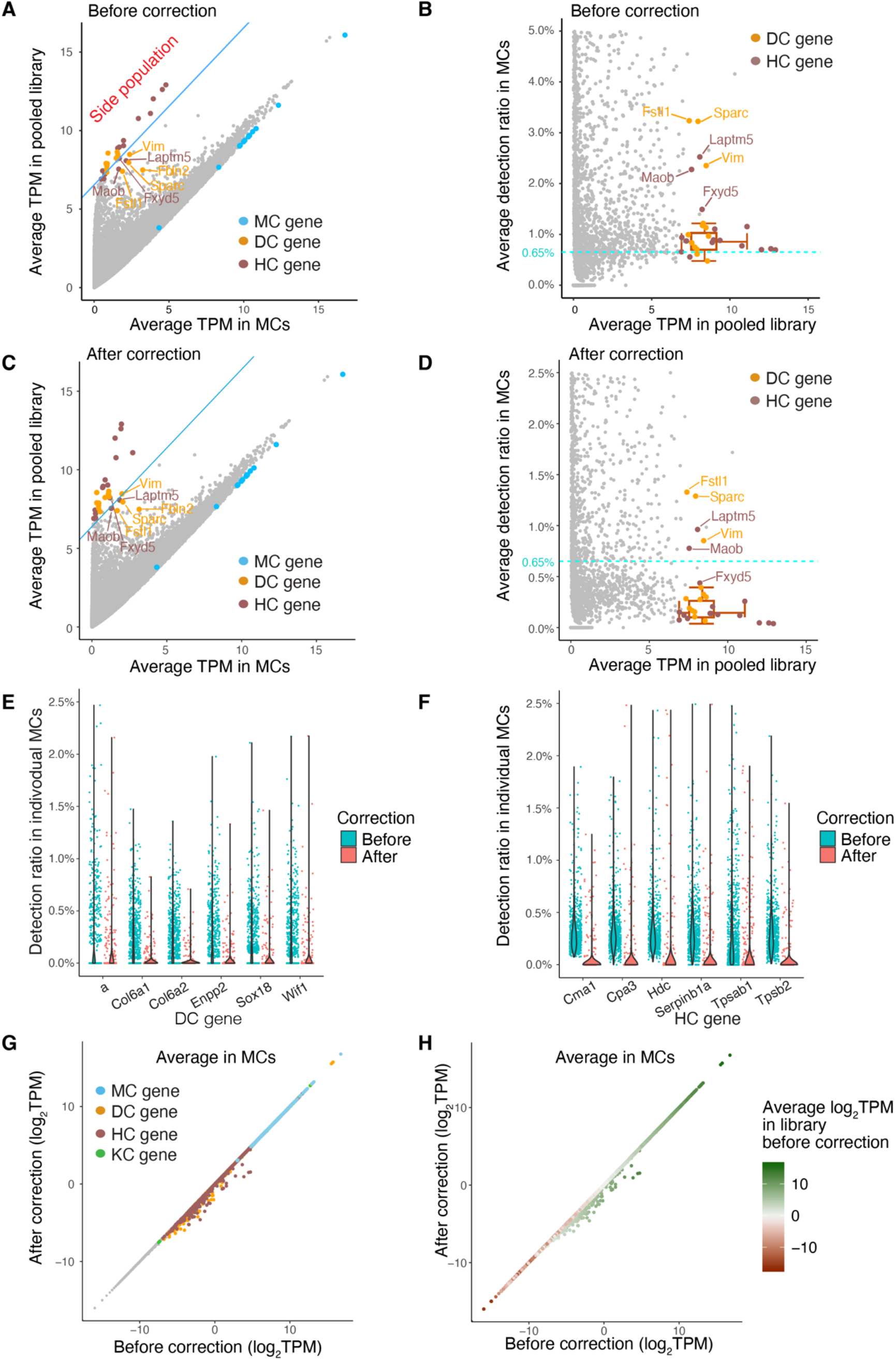
Index hopping correction reduces the detection of non-self marker genes. **(A)** and **(B)** Same as Figure 4A and B – precorrection scatter plots for comparison. **(C)** and **(D)** Scatter plots showing the average TPM (C) and detection ratios (D) of individual genes in single MCs after index hopping correction. Colored labels indicate the top marker genes for different cell types. The blue line in (C) is same as the one in (A) which separates the side population. The cyan dashed line in (B and D) represents the estimated index hopping rate before correction (0.65%). TPMs are displayed on a log2 scale (log2 (average TPM + 1)). Box plots depict the median and interquartile range ± SD of average detection ratio for the side population genes after correction. **(E)** and **(F)** Violin plots showing the detection ratios of the top 6 DC (E) and top 6 HC (F) marker genes in MCs before and after correction. **(G)** and **(H)** Scatter plots showing the log2 scaled average TPM of individual genes in single MCs before and after correction. Colors indicate the marker genes in different cell types **(G)** and represent levels of average TPM in library (H).

Notably, genes in the side population of top HC and DC genes were ranked well above the inflection point, and the non-specific marker genes ranked near the inflection point (Fig. 4D). Consistent with MC and KC having shared epithelial progenitors and similar transcriptomes, the detection ratio of KC marker genes in MCs was typically higher than the index hopping rate reflecting the relative lack of specific KC marker genes (Fig. S5A, B). Following this pattern, in cases where marker genes appeared to be specific to the two cell types being compared, a distinct side population was observed when visualizing average expression levels and detection ratios (Fig. S5C-H).

To address the background gene detection resulting from index hopping, we implemented a simple correction method. We subtracted 0.65% of the average TPM for each gene in our library from the single-cell gene TPM matrix. Following the correction, the detection ratios of non-self genes in MCs (Fig. 5A-F) and other cell types (Fig. S5A’- H’) was notably reduced. Additionally, we investigated the impact of the correction on individual gene TPM values in MCs. As expected, highly expressed HC and DC marker genes showed notably reduced detection in MCs after correction, whereas the reduction in MC marker genes in MCs was inconsequential (Fig. 5G-H, S6A-D). Similarly, when applying the correction in HCs, only non-self genes with high library TPM values were notably impacted, whereas HC marker genes remained comparable to pre-correction levels in HCs (Fig. S6E-J).

### Correcting for index hopping impacts single-cell analysis and interpretation

To assess the impact of index hopping correction on the bioinformatic analysis of the scRNA-seq data, we analyzed our data set with and without correction. To evaluate the heterogeneity among single cells, we employed cell-to-cell distance, which quantifies transcriptomic similarities between cells. A smaller cell distance indicates a higher level of similarity between two cells [15]. We compared the cell distances among all cells in the library before and after correction. As misassigned read contamination would be expected to artificially make distinct cell types more similar, we anticipated that index hopping correction would increase distances between cells. The density plot of cell distances demonstrated that index hopping correction did indeed increase measured heterogeneity within the DC and HC clusters (Fig. 6A, B). This suggests that index hopping correction can mitigate artificial cell-cell similarities caused by randomly misassigned reads across scRNA-seq libraries.

**Figure 6.**
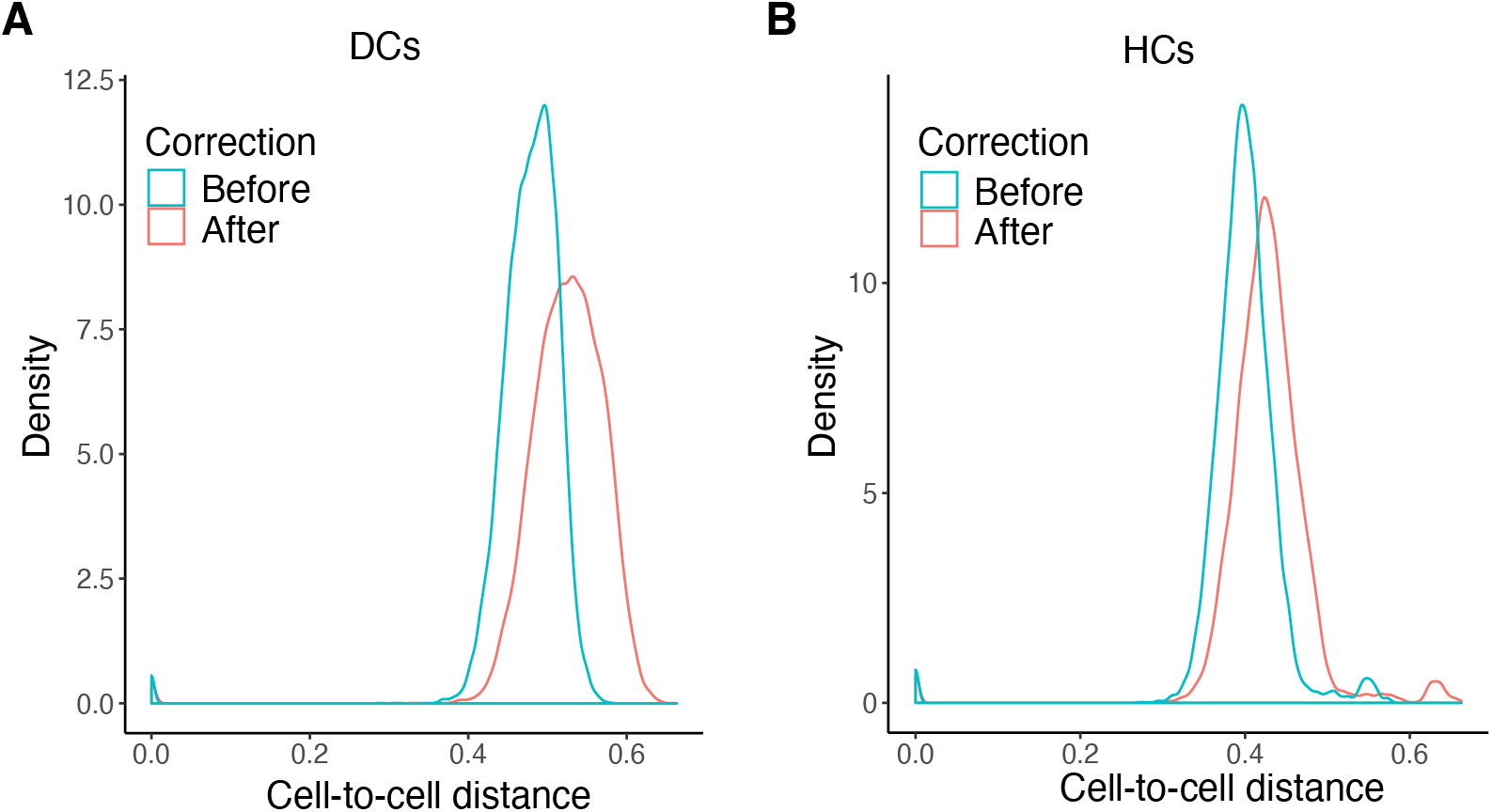
Index hopping correction increases the detection of single cell heterogeneity. Density plots showing the distribution of cell-to-cell distance among individual DCs (A) and HCs (B) before and after correction. Kolmogorov-Smirnov test: p &lt; 2.2 x 10^-16^ for cell-to-cell distance before correction vs. after correction.

Next, we investigated how index hopping correction impacted the analysis of developmental stage classification. Our scRNA-seq dataset included single cells spanning a wide range of developmental timepoints, from embryonic to postnatal skin. Focusing on DCs, we subjected uncorrected and corrected data to Seurat clustering identifying three cell clusters (Fig. S7A, E). Consistent with the cells’ chronological age, each cluster represented a distinct stage of skin development (Fig. S7B, F). We then used Monocle trajectory analysis to plot the single DCs along a developmental pseudotime before (Fig. S7C, D) and after (Fig. S7G, H) index hopping correction.

Comparing the developmental stage classification of each DC before and after correction, we observed that 79.9% of DC cells retained their stage classification whereas 20.1% of cells were reclassified after correction (Fig. 7A). Consistent with stage reclassifications, there was a reordering of the DC pseudotime trajectory after index hopping correction (Fig. 7B), suggesting that index hopping misassignments had impacted the results of both cell clustering and state hierarchy analyses.

**Figure 7.**
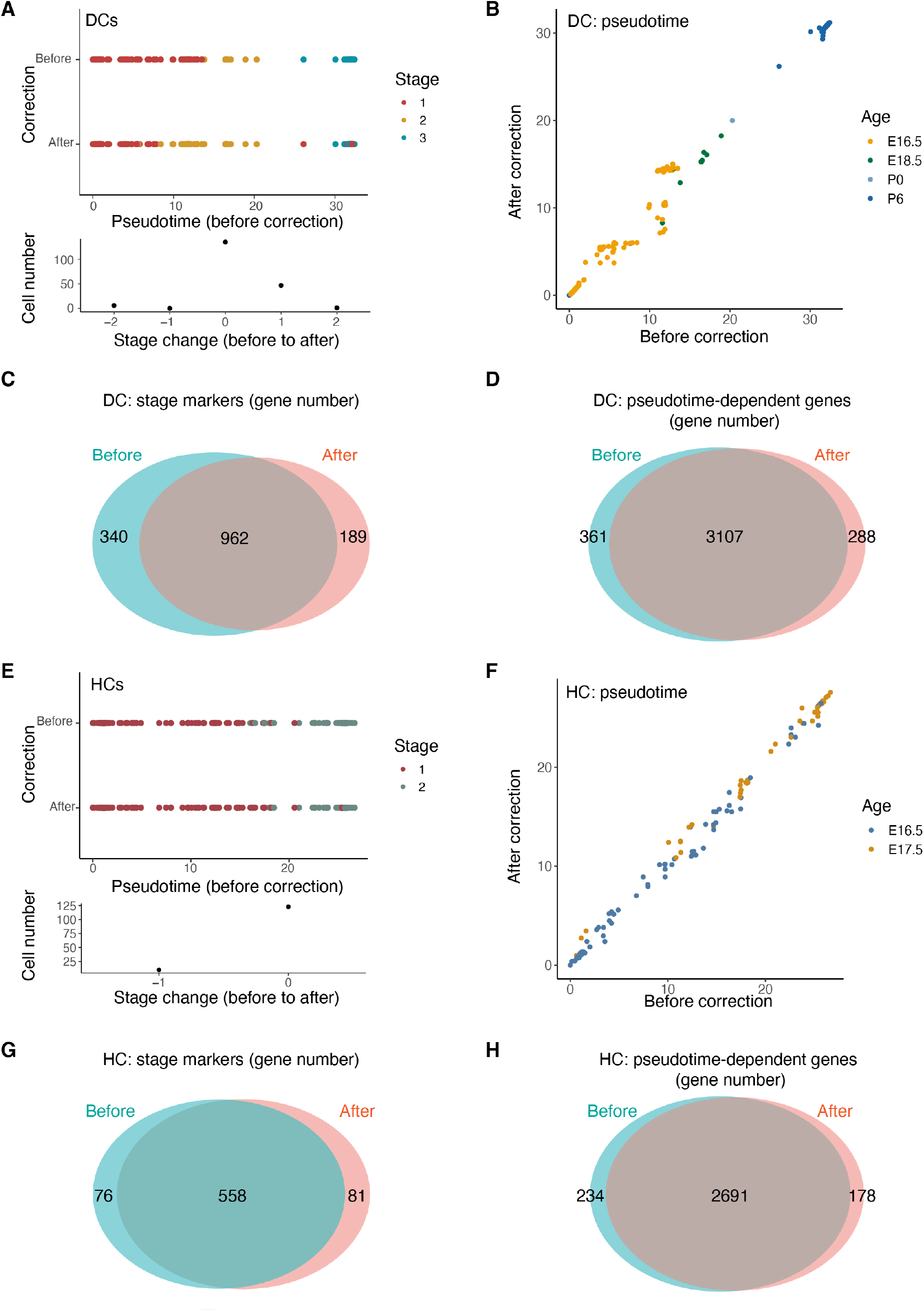
Index hopping correction impacts the single-cell analysis of cell developmental stages. **(A)** and **(E)** Upper: Scatter plots showing the cell developmental stages of individual DCs (A) and HCs (E) assigned by Seurat clustering before and after index hopping correction. Lower: Histogram showing the frequency of cell stage assignment changes from before correction to after correction. **(B)** and **(F)** Scatter plots showing the pseudotime order of individual DCs (B) and HCs (F) identified by Monocle trajectory analysis before and after correction. Colors indicate the age of mouse source for individual cells. **(C)** and (G) Venn diagrams comparing differentially expressed marker genes for any developmental stage of DCs (C) and HCs (G) identified by Seurat before and after correction. **(D)** and **(H)** Venn diagrams comparing pseudotime-dependent genes identified by Monocle for DCs (D) and HCs (H) before and after correction.

We performed a similar pre and post correction comparison for two age groups of HCs (E16.5 and E17.5). Clustering identified two distinct cell clusters before (Fig. S8A, B) and after (Fig. S8E, F) correction, and trajectory analysis revealed a potential developmental trajectory before (Fig. S8C, D) and after (Fig. S8G, H) correction. Index hopping correction also exhibited an analogous, albeit smaller, effect on the stage classification and pseudotime order of some HCs (Fig. 7E, F).

Analysis of differentially expressed genes is often used to define transcript signatures for cell clusters whereas identification of genes that are dynamically expressed over a pseudotime trajectory can suggest biologically relevant genes. Accordingly, we explored the impact of index hopping correction on these analyses. Using Seurat and Monocle, we identified differentially expressed developmental stage marker genes as well as genes that significantly changed over the developmental pseudotime for both DCs and HCs. Comparing the aggregate set of stage marker genes and pseudotime-dependent genes before and after index hopping correction, we observed a substantial overlap in each gene set for DCs (Fig. 7C, D) and HCs (Fig. 7G, H). Nonetheless, index hopping correction altered the composition of these gene lists in each case, once again suggesting that index hopping misassignments can impact the gene composition of cell state signatures. Collectively, these results demonstrate that index hopping measurably influences the biological interpretation of scRNA-seq data. **Hiseq2500 generates a lower index hopping rate compared to Hiseq3000**

As sequencing technology has advanced, the use of patterned flow cells has increased throughput and efficiency. However, sequencing platforms utilizing a patterned flow cell with Exclusion Amplification chemistry, such as the HiSeq3000 and NovaSeq, appear to exhibit higher levels of index hopping compared to platforms without patterned flow cells [3–5, 8, 9]. Our libraries were sequenced using the HiSeq3000 platform. To test if the patterned flow cell technology impacted the degree of index hopping, we re-sequenced one library (Pool #2) on the HiSeq2500, which has a similar sequencing depth but does not employ a patterned flow cell. We compared the gene detection ratios in MCs between the HiSeq2500 and HiSeq3000 platforms. As expected, when analyzing MCs, the detection ratios of genes in the side population, including top HC markers, was reduced on the HiSeq2500 compared to the HiSeq3000, despite the average library TPM remaining unchanged between the two platforms (Fig. 8A-D). The detection ratio of top DC and HC genes in individual MCs was also lower on the HiSeq2500 compared to the HiSeq3000 (Fig. 8E, F). Interestingly, the detection ratio of non-self genes in MCs on the HiSeq2500 was comparable to their HiSeq3000 post-correction levels (Fig. 8A-F, 5C-F, S5A’-H’). Similar trends were observed in HCs, where the detection ratios of non-HC genes were lower on the HiSeq2500 compared to the HiSeq3000 (Fig. S9).

**Figure 8.**
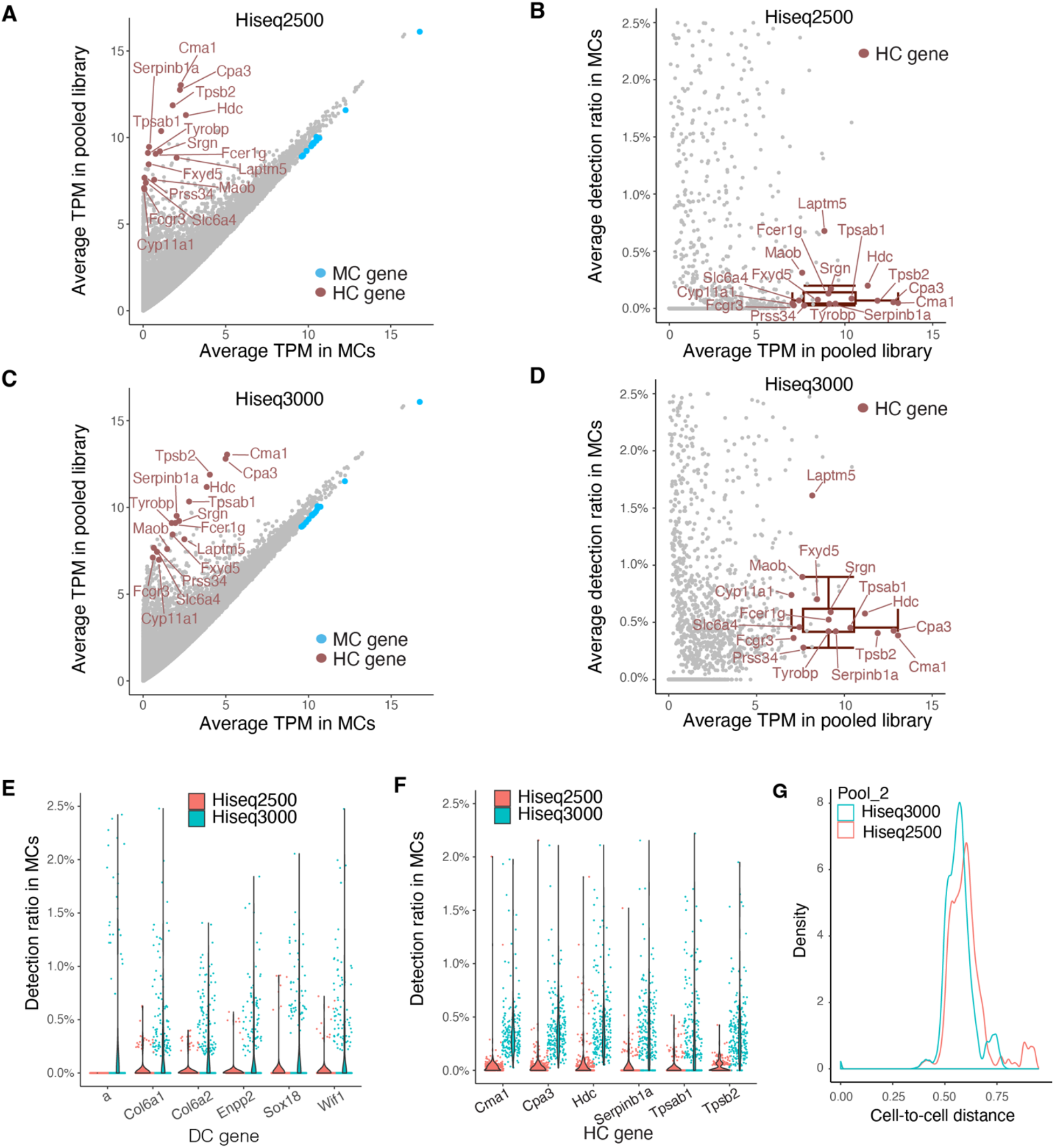
Index hopping is less pronounced in Hiseq2500 platform compared to Hiseq3000. **(A-D)** Scatter plots showing the average TPM (A, C) and detection ratio (B, D) without index hopping correction of individual genes in single MCs in library pool #2 sequenced on Hiseq2500 (A, B) and Hiseq3000 (C, D) platforms. TPMs are displayed on a log2 scale (log2 (average TPM + 1)). Colors indicate the top marker genes for different cell types. Box plots depict the median and interquartile range ± SD of average detection ratio for the color-labeled HC genes. **(E)** and **(F)** Violin plots showing the detection ratios in MCs of the top 6 DC (E) and HC (F) marker genes using sequencing data from Hiseq2500 and Hiseq3000 platforms without index hopping correction. **(G)** Density plot showing the distribution of cell-to-cell distance among single cells in library pool #2 sequenced on Hiseq2500 and Hiseq3000. Kolmogorov-Smirnov test: p &lt; 2.2 x 10^-16^ for cell-to-cell distance from Hiseq2500 vs. Hiseq3000.

Furthermore, the cell distance among single cells in Pool #2 was greater on the HiSeq2500 than on the HiSeq3000 (Fig. 8G). These findings confirm that patterned flow cell technologies are more susceptible to index hopping [3–5, 8, 9], and that our correction strategy compensates for this vulnerability.

## DISCUSSION

In this study, we developed an approach to estimate and correct for index hopping in multiplexed scRNA-seq data. We performed plate-based single cell sequencing of four distinct cell types sorted from mouse skin with a focus on MCs as the most abundant cell type in the libraries. To estimate the rate of read misassignments, we analyzed differentially expressed genes across each cell type. Top genes expected to be expressed exclusively in their respective cell types were designated as marker genes, and the presence of non-self marker genes in a particular cell was interpreted as contamination from index hopping. We calculated the gene detection ratio for each gene in MCs and estimated the index hopping rate based on the ratio of non-self genes.

Since the detection ratio of non-self genes in MCs was consistent regardless of their overall detection levels in the library, we assumed a fixed index hopping rate across all samples in the libraries. To correct for index hopping, we reduced the TPM for each gene by the index hopping rate multiplied by the transcript’s average expression across the library. The correction significantly reduced read misassignments, as evidenced by the decreased detection ratio of non-self genes. Comparing data obtained from Hiseq3000 and Hiseq2500 sequencing platforms confirmed that index hopping was more pronounced in platforms employing patterned flow cells (e.g., Hiseq3000/4000/X, NextSeq 1000/2000, NovaSeq X/X Plus, and NovaSeq 6000). Furthermore, we evaluated the impact of index hopping correction on biological analysis. Our findings revealed that correction of index hopping mitigated artificial cell-cell similarities, impacted the results of cell clustering and state hierarchy analyses, and influenced the gene composition of cell state signatures. In summary, we propose a simple and practical methodology for estimating and correcting index hopping that appears to improve the accuracy of transcriptome data and the cell biology inferred from its analysis.

By strategically pooling cells with distinct transcriptomes, we were able to effectively identify and characterize the extent of index hopping. We propose that this method can be adapted to detect index hopping in multiplex sequencing experiments involving both heterogeneous and homogeneous libraries. In the case of a homogeneous cell population, one approach is to introduce single cells from a different cell type as "spike-ins" during the library preparation process. This deliberate addition of distinct cells will introduce non-self genes into the otherwise homogenous cell library allowing for the generation of a side population of non-self genes if index hopping events occur. By leveraging this spike-in strategy, researchers can effectively assess the presence and impact of index hopping contamination, thus enhancing the quality and reliability of scRNA-seq data in various experimental settings.

Our approach to index hopping has certain limitations. An Illumina white paper demonstrated a correlation between the concentration of free index primers and the rate of index hopping [10]. Variations in the amounts of cDNA and free index primers can produce concentration inconsistencies when multiplexing and pooling libraries. To mitigate this variability, we normalized single-cell cDNA concentrations prior to tagmentation and indexing. However, multiplexed libraries with varied cDNA concentrations per sample may result in differing index hopping rate across libraries. Consequently, estimating a universal index hopping rate for correction may not be possible. Therefore, we recommend normalizing cDNA concentration prior to indexing when possible. In addition, our approach was applied to dual indexed multiplexed libraries. Although unique indexing has been proposed as a strategy to address index hopping, this issue can still arise, especially due to the amplification of free index primers on sequencing platforms with patterned flow cells, as evidenced by the droplet-based scRNA-seq studies [4]. Finally, our comparative analyses of scRNA-seq data before and after correction revealed substantial effects on biological analyses. It is crucial to note that the impact of correction on biological analysis may hinge on the specific index hopping rate within each dataset. Balancing the decision to correct for index hopping is paramount, as over-correction can also introduce false results, particularly in analyses that are sensitive to minor changes in gene expression, such as genes with low expression levels. Each experimental scenario requires a thoughtful assessment to determine the most appropriate course of action when considering index hopping correction.

## CONCLUSIONS

Our comprehensive analysis of index hopping in plate-based deep scRNA-seq data suggests that having cells with distinct transcriptomes, due to intrinsic diversity or the inclusion of cell "spike-ins" during library preparation, can serve as a valuable strategy to address index hopping data contamination. To compensate for index hopping, we developed a simplified method that enables both the estimation of the index hopping rate and the subsequent correction of read misassignments. However, it is crucial to consider the specific biological questions being addressed by the dataset when making the decision to apply index hopping correction. By employing our approach, researchers can gain a better understanding of the index hopping phenomenon and its impact on scRNA-seq data. This knowledge can help improve the accuracy and reliability of downstream analyses, ultimately leading to more robust and meaningful biological insights.

## LIST OF ABBREVIATIONS

scRNA-seq: single-cell RNA sequencing
SOX2: SRY-Box Transcription Factor 2
GFI1: Growth Factor Independence 1
MC: Merkel cell
DC: dermal papilla cell
HC: hematopoietic cell
KC: keratinocyte
TPM: transcripts per million

## DECLARATIONS

### Ethics approval

All experiments were performed in accordance with institutional guidelines according to Institutional Animal Care and Use Committee-approved protocols.

## Authors’ contributions

Conceptualization: IB; Methodology: LM, SD, MCK; Data curation: LM, SD; Resources: IB, SD; Formal analysis: LM, JJ, JH; Investigation: LM, LC, SB, YS; Writing - original draft preparation: LM; Writing - review and editing: all authors; Supervision: IB, HK; Funding acquisition: IB.

## Acknowledgements

We thank Drs. Livak Ferenc and Teresa Hawley from the CCR Flow Cytometry Core, as well as Drs. James Simone, Jeff Lay and Kevin Tinsley from the NIAMS Flow Cytometry Section, for their assistance with cell sorting. We acknowledge Dr. Valery Bliskovsky from the CCR Genomics Core for help with single-cell cDNA library preparation, Dr. Sean Conlan for scientific discussion, and Faiza Naz from the NIMAS Genomic Technology Section for her assistance with sequencing. This work utilized the computational resources of the NIH HPC Biowulf cluster (https://hpc.nih.gov).

## SUPPLEMENTARY MATERIAL

Figures S1-S9:

Figure S1. Skin schematic showing cells labeled by GFP in Sox2GFP/+ and Gfi1GFP/+ reporter mice.

Figure S2. Index hopping can be detected when occurring among different cell types. **Figure S3.** Top DC markers genes are detected in MCs with single index hopping potential from DCs and the detection ratio is correlated with number of different cell types with the potential for index hopping.

Figure S4. Rank ordered scatter plot showing distribution of gene detection ratios in MCs across all genes from mm10 genome annotated by Ensembl.

Figure S5. Index hopping correction reduces the detection of non-self marker genes in different cell types.

Figure S6. Index hopping correction reduces the detection of non-self marker genes. **Figure S7.** Assigned developmental stages of single DCs are altered by index hoping correction.

Figure S8. Assigned developmental stages of single HCs are altered by index hoping correction.

Figure S9. Detection ratios of top MC marker genes in HCs show that index hopping is reduced on Hiseq2500 compared to Hiseq3000 sequencing platforms.

