## Supplementary material for "Estimating and correcting index hopping misassignments in single-cell RNA-seq data"

### Figures S1-S9

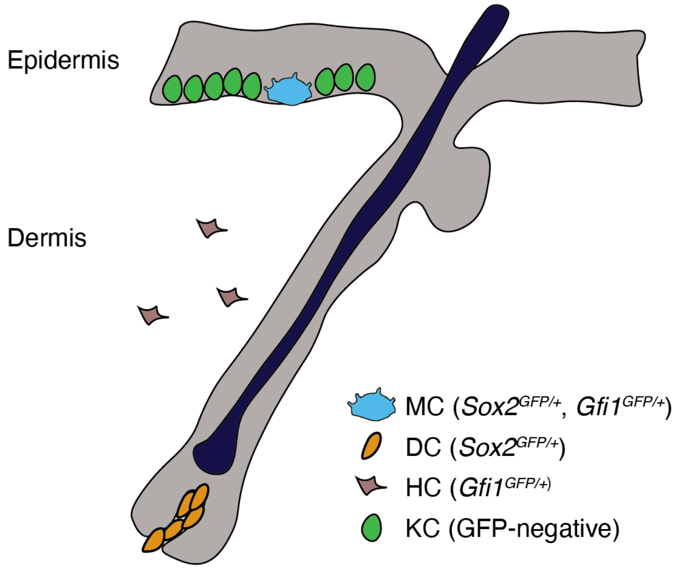


**Figure S1. Skin schematic showing cells labeled by GFP in *Sox2$^{GFP/+}$* and *Gfi1$^{GFP/+}$* reporter mice.** GFP expresses in Merkel cells (MCs) in the *Sox2$^{GFP/+}$* and *Gfi1$^{GFP/+}$* epidermis. GFP expresses in dermal papilla cells (DCs) in the *Sox2$^{GFP/+}$* dermis. GFP expresses in intravascular and tissue-resident hematopoietic cells (HCs) of *Gfi1$^{GFP/+}$* mice.

A

B

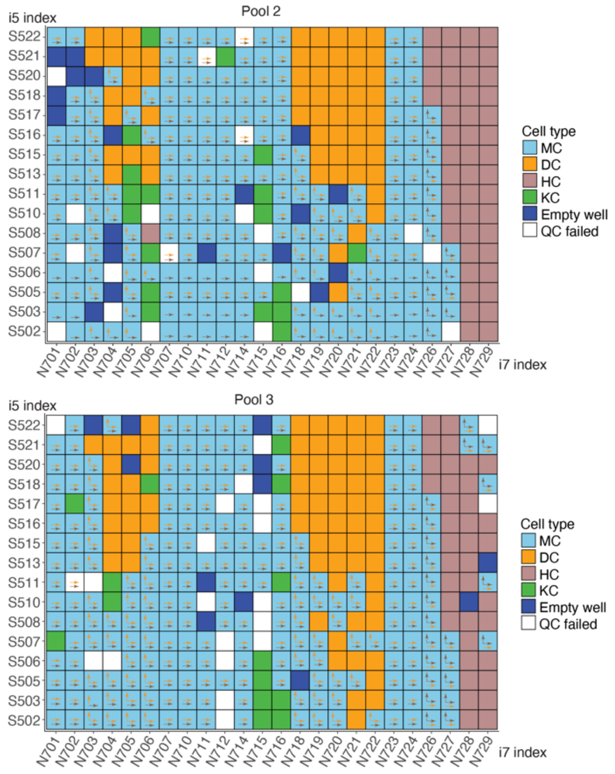


C

| | MC | DC | HC | KC | Empty well | QC failed |
|---|---|---|---|---|---|---|
| Pool 1 | 206 | 58 | 50 | 47 | 6 | 17 |
| Pool 2 | 219 | 60 | 48 | 18 | 19 | 20 |
| Pool 3 | 229 | 71 | 37 | 14 | 12 | 21 |
| Total | 654 | 189 | 135 | 79 | 37 | 58 |

D

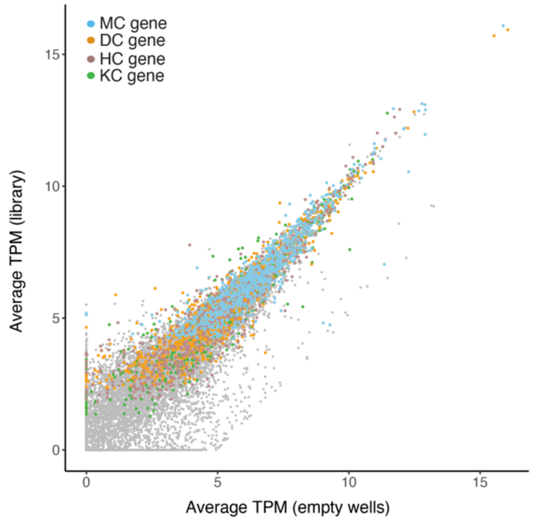


**Figure S2. Index hopping can be detected when occurring among different cell types.**
**(A)** and **(B)** Overview of the indexing used for 384 single-cell cDNA libraries in the second and third pooled sequencing libraries. Cell types, along with empty wells and quality control (QC) failed wells are color-coded. Arrows show for each individual MCs, the potential index hopping with DCs (orange arrows) or HCs (brown arrows). Vertical arrows show i7 index hopping, and horizontal arrows show i5 index hopping.
**(C)** Table showing the number of single cells for different cell types, empty wells and QC failed wells in each pooled sequencing library.
**(D)** Scatter plot showing the correlation of average TPM of all genes in empty wells and the entire library, with values transformed using a log2 scale (log2(average TPM +1). Colors indicate marker genes for the 4 cell types as shown in Figure 2B.

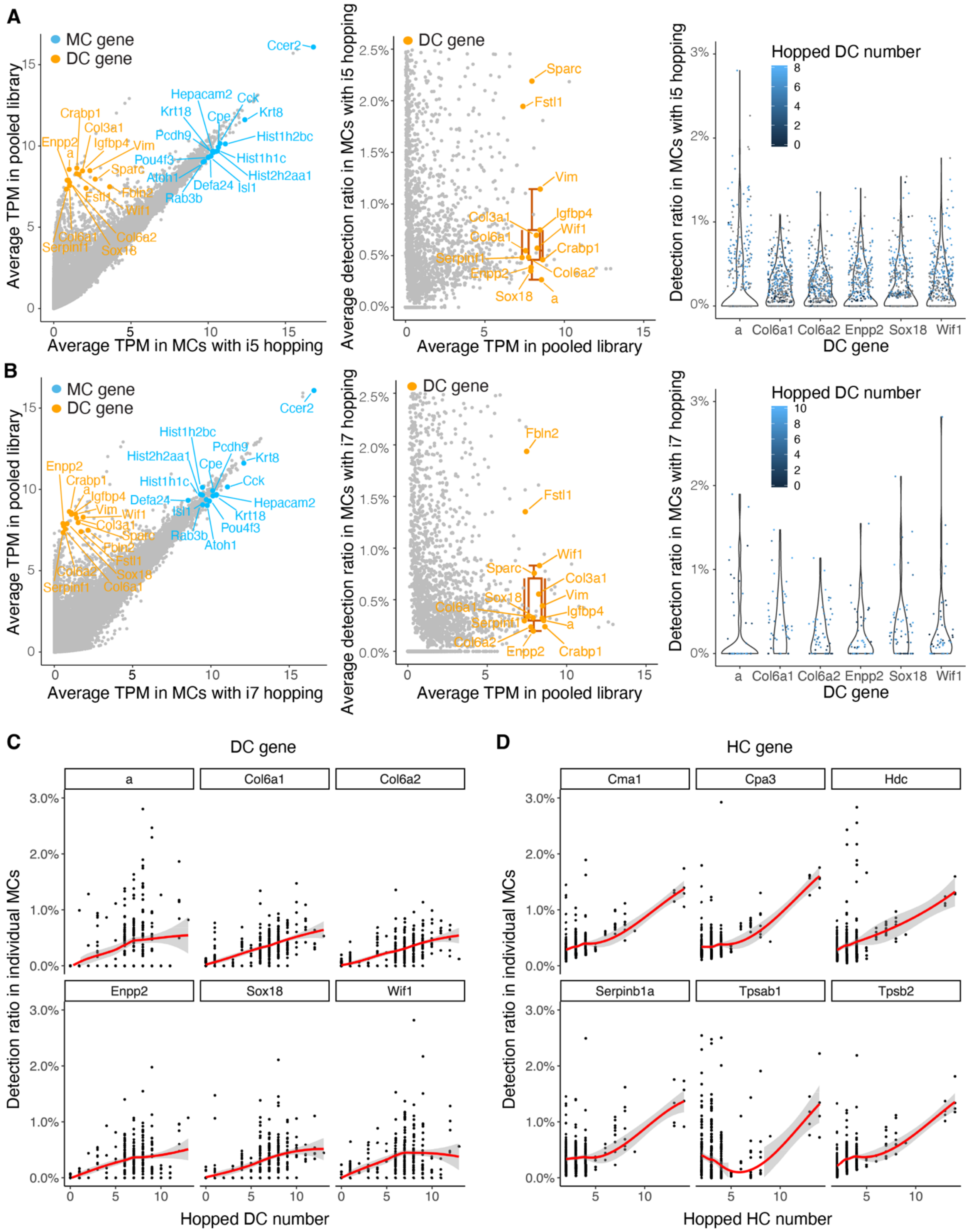
A
B
C
D
MC gene
DC gene
Hopped DC number
Average TPM in pooled library
Average TPM in MCs with i5 hopping
Average TPM in MCs with i7 hopping
Average detection ratio in MCs with i5 hopping
Average detection ratio in MCs with i7 hopping
Detection ratio in MCs with i5 hopping
Detection ratio in MCs with i7 hopping
Detection ratio in individual MCs
Hopped HC number
HC gene
a
Col6a1
Col6a2
Enpp2
Sox18
Wif1
Cma1
Cpa3
Hdc
Serpinb1a
Tpsab1
Tpsb2
Ccer2
Hepacam2
Cck
Krt18
Cpe
Krt8
Pcdh9
Hist1h2bc
Hist1h1c
Pou4f3
Hist2h2aa1
Atoh1
Isl1
Defa24
Rab3b
Crabp1
Col3a1
Igfbp4
Vim
Sparc
Fbln2
Fstl1
Serpinf1

**Figure S3. Top DC markers genes are detected in MCs with single index hopping potential from DCs and the detection ratio is correlated with number of different cell types with the potential for index hopping.**
MCs had i5 **(A)** or i7 **(B)** index hopping potential from DCs: Left, scatter plots showing the average TPM of individual genes in single MCs correlated to their average TPM in the library. Middle, scatter plots showing the average detection ratio of individual genes in single MCs. TPMs are displayed on a log2 scale (log2 (average TPM + 1)). Colored labels indicate top marker genes for different cell types. Box plots show the median and interquartile range ± SD of average detection ratio for color-labeled DC genes. Right, violin plots showing the detection ratio of the top 6 DC marker genes in individual MCs.
**(C)** Scatter plot showing detection ratios of the top 6 DC marker genes in individual MCs correlated with the number of DCs with potential for index hopping.
**(D)** Scatter plot showing detection ratios of the top 6 HC marker genes in MCs correlated with the number of HCs with the potential for index hopping.

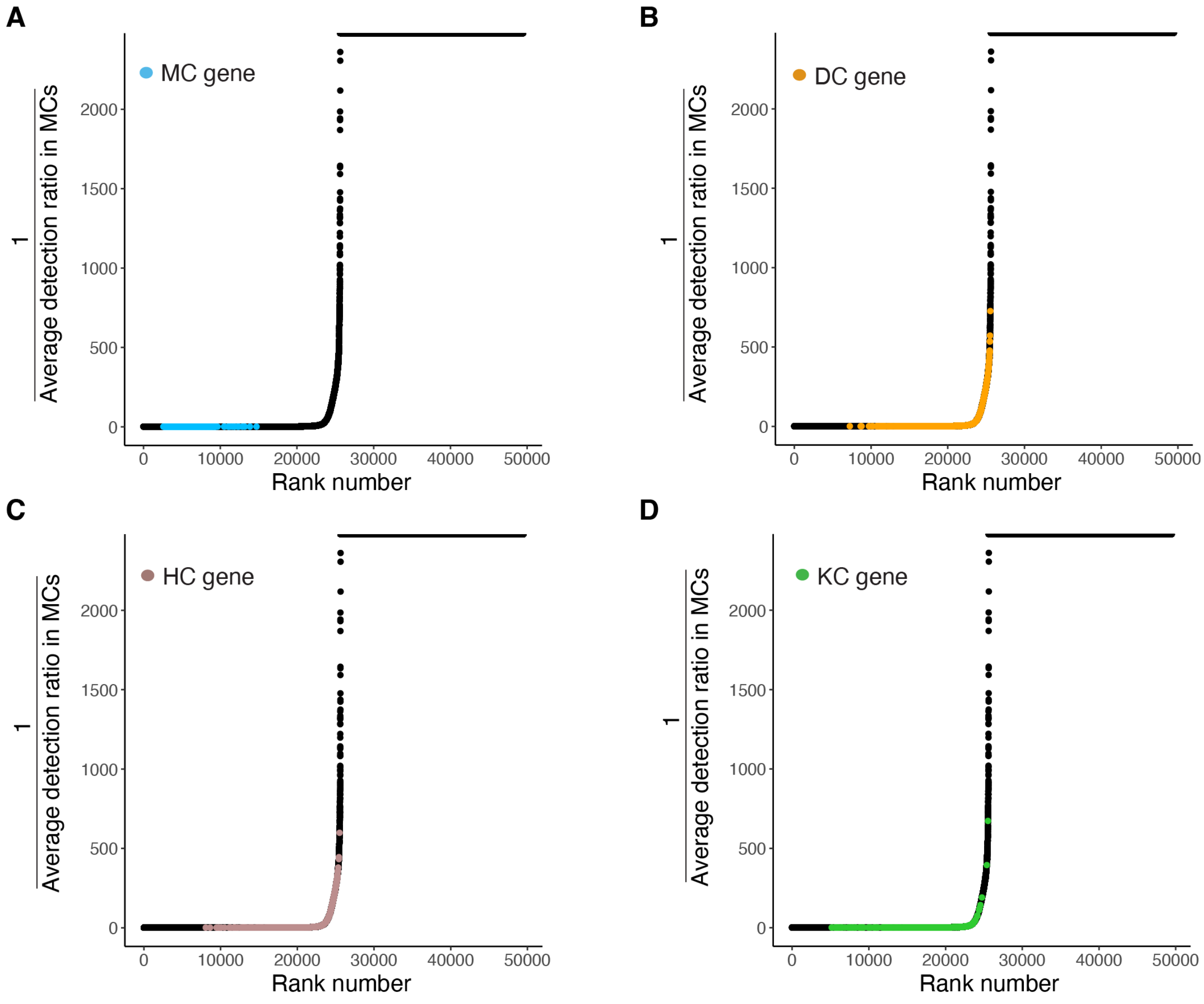


**Figure S4. Rank ordered scatter plot showing distribution of gene detection ratios in MCs across all genes from mm10 genome annotated by Ensembl.** MC **(A)**, DC **(B)**, HC **(C)**, and KC **(D)** marker genes indicated with different colors.

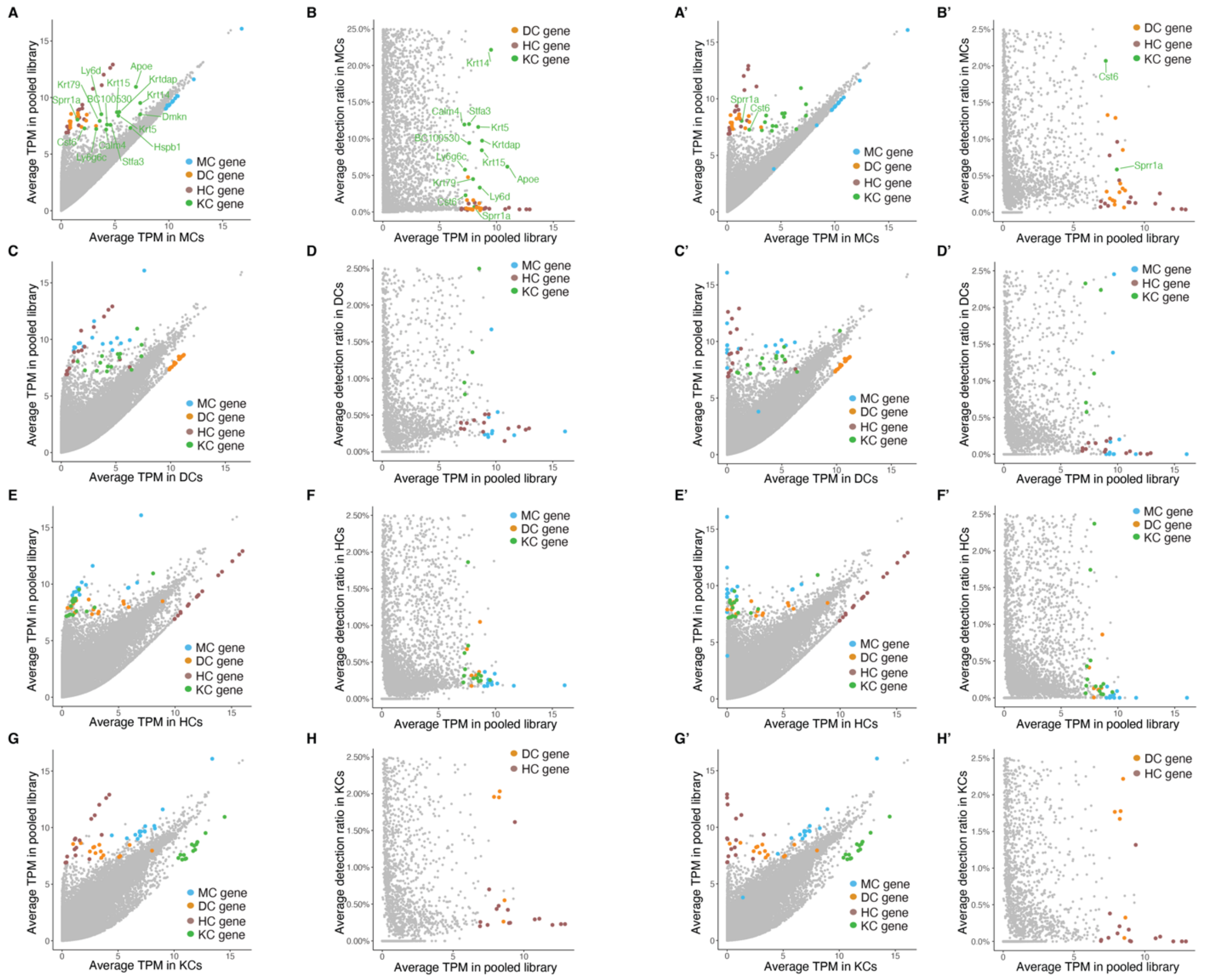

**Figure S5. Index hopping correction reduces the detection of non-self marker genes in different cell types.**
**(A-H),** before correction. **(A'-H'),** after correction. **(A, C, E, G, A', C', E', G')** Scatter plots showing the average TPM of individual genes in single MCs (A, A'), DCs (C, C'), HCs (E, E'), and KCs (G, G') before and after correction. Marker genes for different cell types are indicated with colors. **(B, D, F, H, B', D', F', H)** Scatter plots showing the average gene detection ratios of individual genes in single MCs (B, B'), DCs (D, D'), HCs (F, F'), and KCs (H, H'). TPMs are displayed on a log2 scale (log2 (average TPM + 1)). Marker genes for different cell types are indicated by color or labeled.

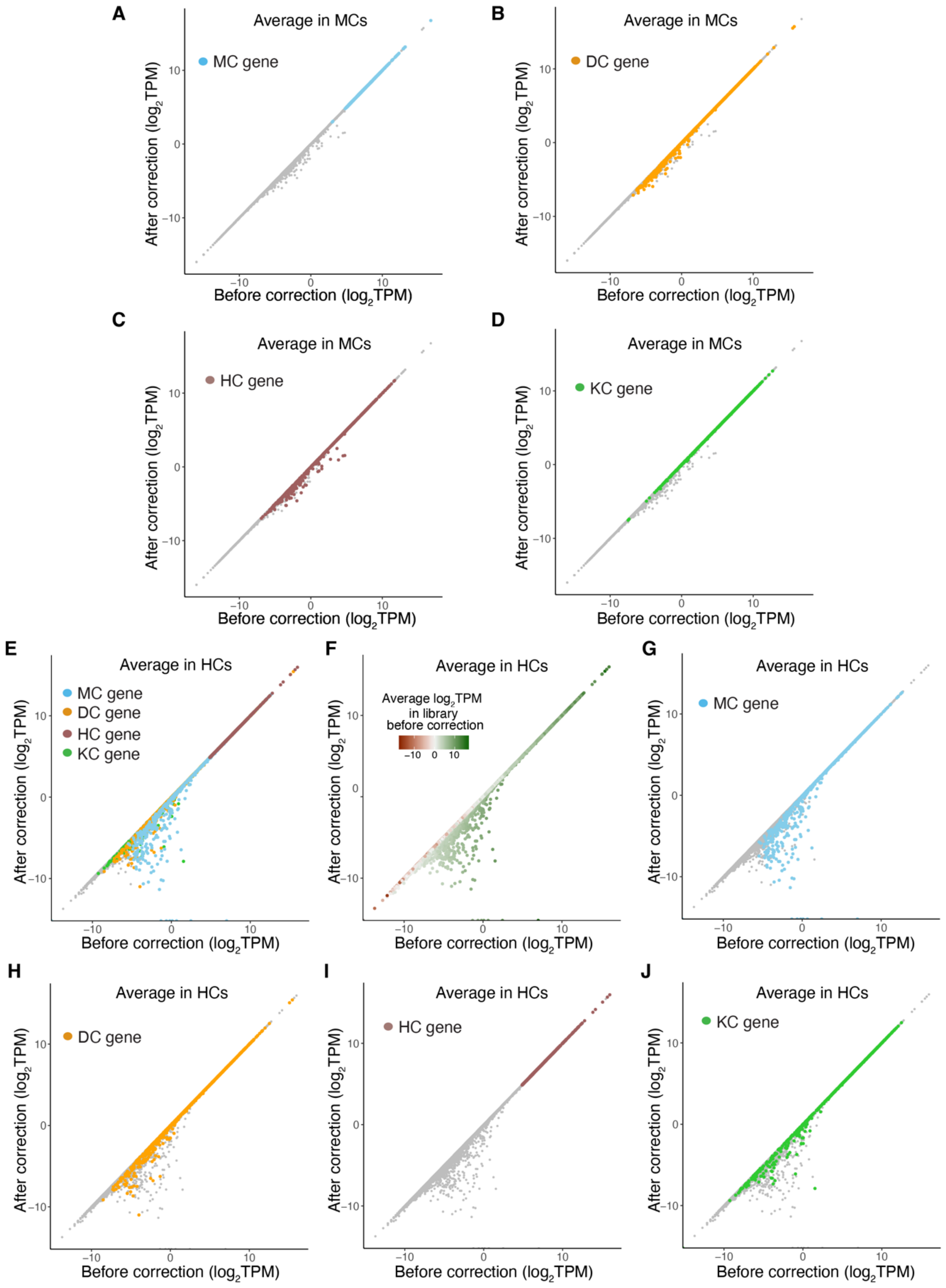


**Figure S6. Index hopping correction reduces the detection of non-self marker genes.**
Scatter plot showing the log2 scaled average TPM of individual genes in single MCs **(A-D)** and HCs **(E-J)** before and after correction. Colors indicate the marker genes for different cell types **(A-E, G-J)** and represent average TPM in library **(F)**.

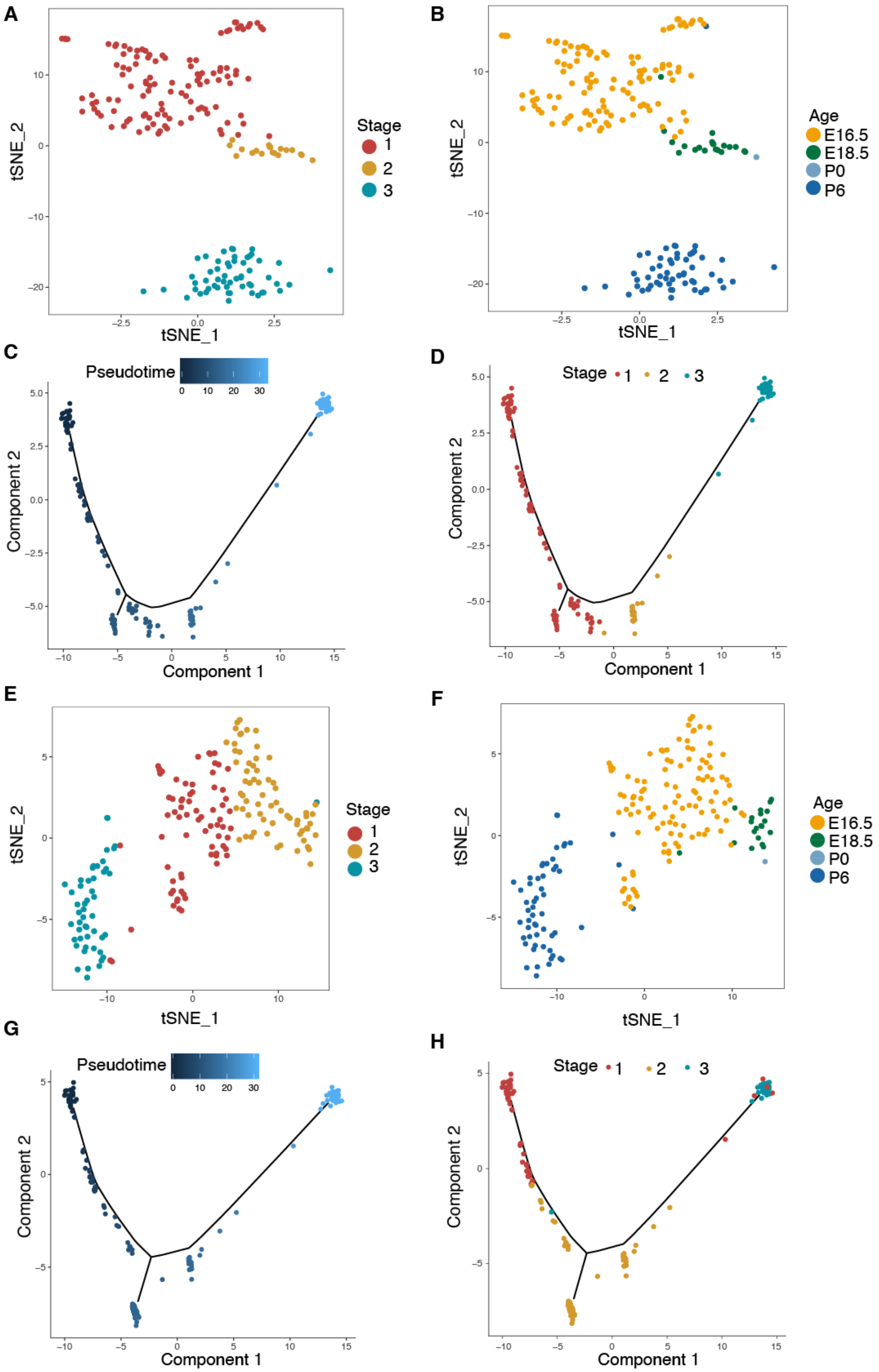


**Figure S7. Assigned developmental stages of single DCs are altered by index hoping correction. (A, B)** and **(E, F)** tSNE plots showing clustering of the gene expression profiles of single DCs before (A, B) and after (E, F) correction. Colors indicate cell developmental stages identified (A, E) or age of source mouse for individual cells (B, F).

**(C, D)** and **(G, H)** Trajectory plots showing developmental pseudotime trace of single DCs before (C, D) and after (G, H) correction. Colors indicate pseudotime identified by Monocle (C, G) and developmental stage identified by Seurat (D, H) for each cell.

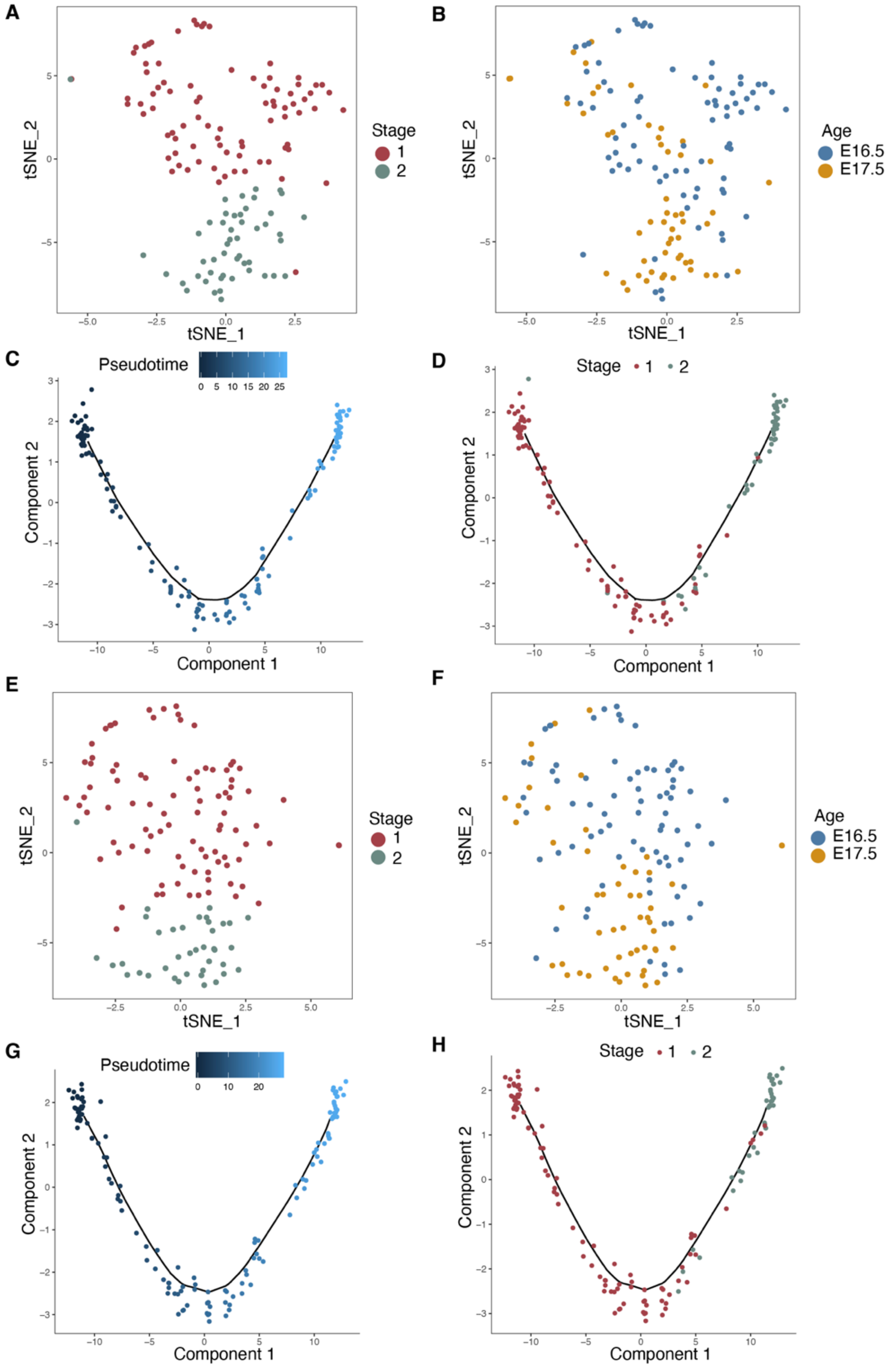

**Figure S8. Assigned developmental stages of single HCs are altered by index hoping correction.**
**(A, B)** and **(E, F)** tSNE plots showing clustering of the gene expression profiles of single HCs before (A, B) and after (E, F) correction. Colors indicate cell developmental stages identified (A, E) or age of source mouse for individual cells (B, F).
**(C, D)** and **(G, H)** Trajectory plots showing developmental pseudotime trace of single HCs before (C, D) and after (G, H) correction. Colors indicate pseudotime identified by Monocle (C, G) and developmental stage identified by Seurat (D, H) for each cell.

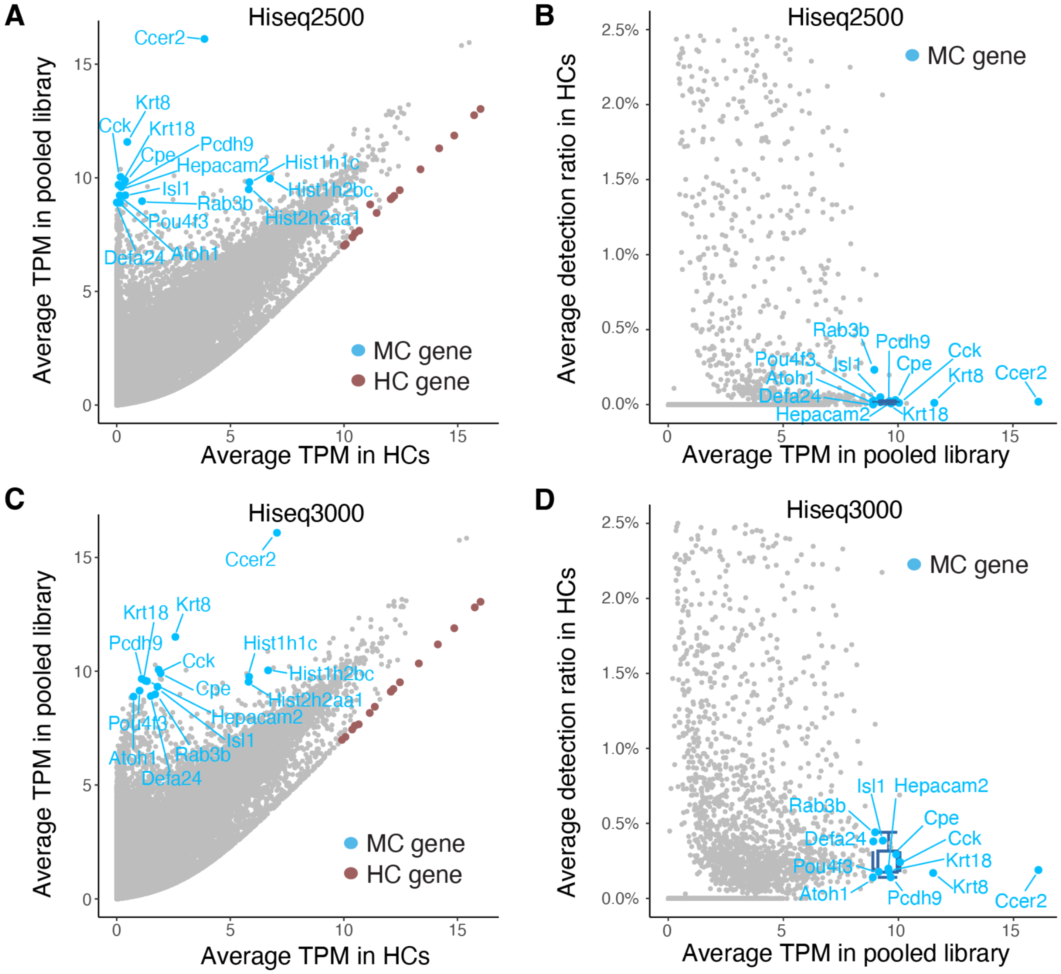


**Figure S9. Detection ratios of top MC marker genes in HCs show that index hopping is reduced on Hiseq2500 compared to Hiseq3000 sequencing platforms.**
Scatter plots showing the average TPM (A, C) and detection ratio (B, D) of individual genes in single HCs sequenced on Hiseq2500 (A, B) and Hiseq3000 (C, D) platforms. TPMs are displayed on a log2 scale (log2 (average TPM + 1)). Colored labels show marker genes for different cell types. Box plots show the median and interquartile range ± SD of average detection ratio for color-labeled MC genes in HCs.
